# Transposable elements shape olaparib response according to BRCA1 status in triple-negative breast cancer

**DOI:** 10.64898/2026.07.15.738694

**Authors:** Daniela Moreira Mombach, Carlos Mendez-Dorantes, Rafael L. V. Mercuri, Phillip Schofield, Suelen Cristina Soares Baal, Maria Augusta Poersch, Kathleen H. Burns, Jaqueline Carvalho de Oliveira, Elgion Lucio Silva Loreto, Pedro A. F. Galante

**Affiliations:** Programa de Pós-Graduação em Genética e Biologia Molecular, Universidade Federal do Rio Grande do Sul, Porto Alegre, Rio Grande do Sul, Brazil; Hospital Sírio-Libanês, São Paulo, São Paulo, Brazil; Department of Pathology, Dana-Farber Cancer Institute, Boston, Massachusetts 02115, USA; Department of Pathology, Harvard Medical School, Boston, Massachusetts 02115, USA; Broad Institute of Massachusetts Institute of Technology and Harvard, Cambridge, Massachusetts 02142, USA; Departamento de Genética, Universidade Federal do Paraná, Curitiba, Paraná, Brazil; Departamento de Bioquímica e Biologia Molecular, Universidade Federal de Santa Maria, Santa Maria, Rio Grande do Sul, Brazil

**Keywords:** triple-negative breast cancer, LINE-1, transposable element, parp inhibitor, BRCA1, olaparib

## Abstract

**Background:** Triple-negative breast cancer (TNBC) is an aggressive subtype with limited therapeutic options. While PARP inhibitors, such as olaparib, show promise in BRCA1-deficient TNBC through synthetic lethality, up to 50% of patients fail to respond, highlighting the need to understand the molecular mechanisms underlying PARP inhibitors efficacy. Transposable elements (TEs), particularly LINE-1 elements, are increasingly recognized as modulators of genomic instability associated with DNA repair processes and potential key players in synthetic lethality. Here, we investigate the functional relationship between TE activity and olaparib treatment in TNBC with distinct *BRCA1* functional status.

**Methods:** We performed comprehensive multi-OMICs analysis of four TNBC cell lines (two BRCA1-deficient: SUM1315 and MDA-MB-436; two BRCA1-proficient: MDA-MB-468 and BT549) treated with olaparib. We analyzed expression and differential expression of protein-coding genes, TEs, and gene-TE chimeric transcripts. Long-read whole-genome sequencing was employed to detect de novo TE insertions, complemented by a functional assay to quantify LINE-1 retrotransposition activity in olaparib-treated cells.

**Results:** Olaparib treatment induces extensive transcriptomic and genomic disorganization mediated by TEs, especially LINE-1, exclusively in BRCA1-deficient cells. We observed aberrant overexpression of both genes and TEs, including gene–TE chimeric transcripts harboring poison exons within tumorigenic genes and multi-exonic TE–TE chimeras capable of forming immunostimulatory double-stranded RNA (dsRNA) structures. Functional enrichment analyses revealed activation of antiviral immune pathways linked to LINE-1 activity. Consistently, orthogonal assays confirmed LINE-1 retrotransposition in BRCA1-deficient cells following olaparib exposure.

**Conclusions:** Our findings demonstrate that olaparib treatment induces TE activation especially in BRCA1-deficient cells, a novel mechanism that may underlie synthetic lethality in TNBC. This TE activation triggers immune responses and genomic instability, providing new therapeutic opportunities through immunotherapy combinations and suggesting that TE activity may serve as a potential biomarker for treatment stratification of TNBC.

**Graphical Abstract:** 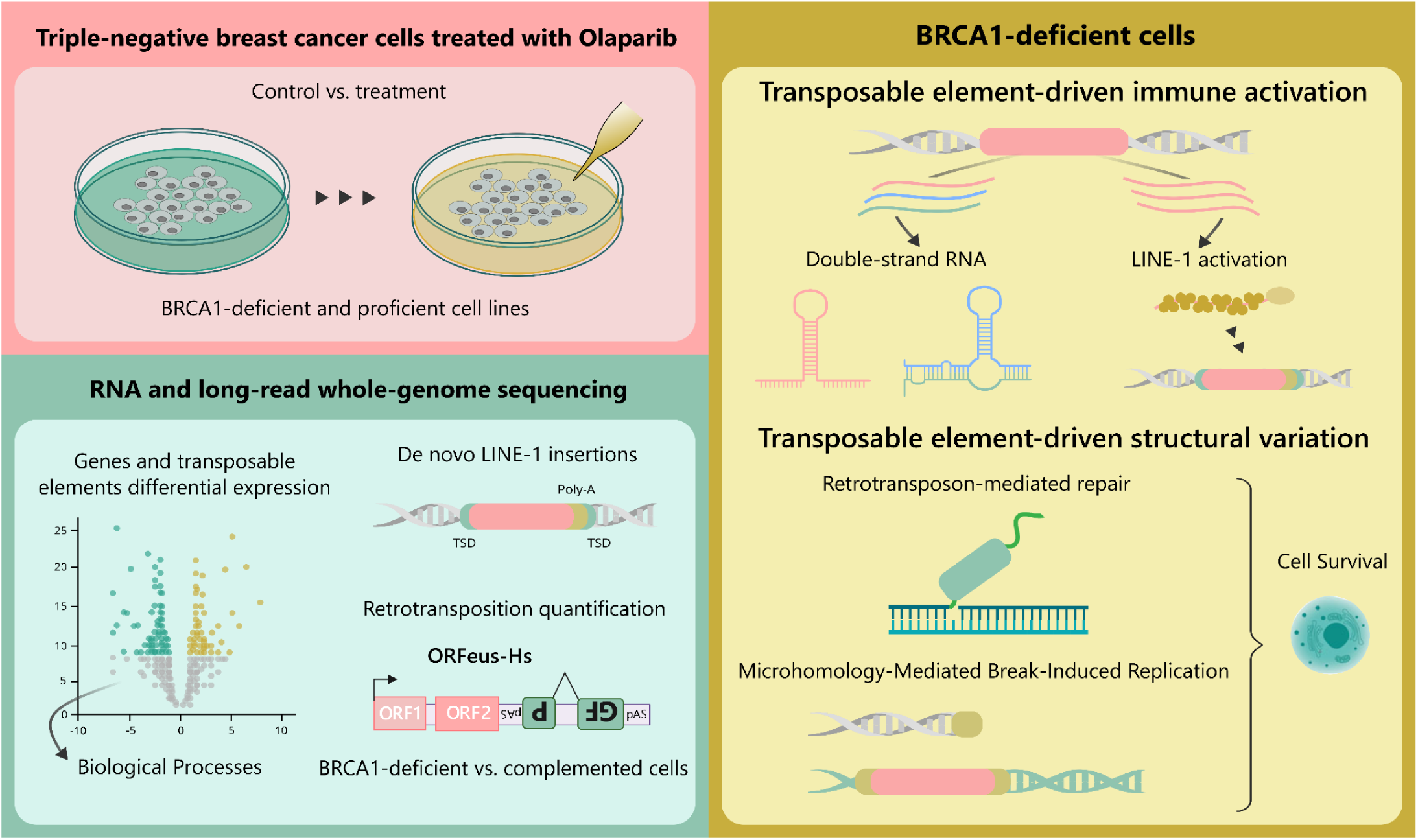

## INTRODUCTION

Breast cancer (BC) remains the most prevalent and the leading cause of cancer-related deaths among women globally, with early-stage, non-metastatic cases often treatable [1]. However, Triple-Negative Breast Cancer (TNBC), an aggressive BC subtype characterized by the absence of estrogen receptor, progesterone receptor, and HER2 expression, presents significant therapeutic challenges due to its high metastatic potential, frequent relapse, and poor prognosis [2,3]. TNBC is frequently associated with *TP53* mutations [3] and up to 20% of patients carry *BRCA1 or BRCA2* (*BRCA1/2*) mutations [4], with higher prevalence in Black and Hispanic populations [5]. Recent advances include PARP inhibition therapy, particularly for BRCA1/2-deficient TNBC, and have shown promise in improving progression-free survival, with drugs like olaparib leveraging synthetic lethality to induce cell death by accumulating irreparable DNA damage in cells with compromised homologous recombination (HR) repair [6]. Despite these therapeutic advances, resistance mechanisms to olaparib treatment lead to disease relapse. Accordingly, the heterogeneous response to PARP inhibitors and tumor recurrences [7–9] highlight the need to identify additional molecular players that may guide this therapeutic strategy.

Transposable elements (TEs) are repetitive DNA sequences with significant regulatory and mutagenic potential [10–12]. In the human genome, Long Interspersed Element-1 human-specific (L1Hs), the only currently active protein-coding element [13], encodes proteins ORF1p [14,15] and ORF2p [16,17], which facilitate their (and other RNAs *in trans*) retrotransposition via target-primed reverse transcription [18], typically resulting in *de novo* insertions flanked by target site duplications [18,19]. In cancer cells, particularly TNBC, epigenetic mechanisms that suppress TE activity, such as L1Hs promoter methylation [20], become disrupted, leading to aberrant TE activation [20]. Somatically acquired LINE-1 insertions are particularly prevalent in tumors with high chromosomal instability, where L1Hs retrotransposition may drive DNA damage [11], genomic deletions [10], and chromosomal rearrangements [12], a process that may be further exacerbated by defective DNA repair due to mutations in genes such as *BRCA1* [21]. Moreover, the frequent co-occurrence of *TP53* mutations in TNBC cells creates a permissive environment for sustained L1Hs expression [22] and genomic instability.

The intricate relationship between DNA repair deficiencies and TE activity becomes especially relevant in TNBC, where *BRCA1* mutations and PARP inhibitor (PARPi) therapy intersect. When functional BRCA1 or BRCA2 (BRCA1/2) is absent, HR-mediated repair of double-strand DNA breaks (DSBs) is compromised and PARPi therapies can cause complex DNA lesions, forcing reliance on other DNA repair pathways, including PARP proteins (PARP1/2) for DNA repair via base-excision repair pathways [23]. While PARPis exploit this vulnerability through synthetic lethality, the substantial treatment failure rate (up to 50% of germline BRCA1/2-deficient patients not responding [7–9]) suggests additional complexity, which may be associated with the genomic instability and abnormal transcriptional profiles.

In this study, we investigate, in TNBC with distinct *BRCA1* mutational status, the functional relationship between the TE activity, BRCA1-deficiency, and olaparib treatment. Through comprehensive transcriptome profiling and long-read whole-genome sequencing (WGS), we systematically analyzed TE activity, expression, and functional consequence in olaparib treatment versus controls cells in both BRCA1-deficient and proficient backgrounds. Our results reveal how the TE activities associated with BRCA1 status fundamentally shape genomic and transcriptomic changes in response to PARP inhibition. Additionally, we provide novel mechanistic insights, based on the TE activities, for the synthetic lethal interaction between BRCA1 deficiency and PARPi treatment in TNBC.

## RESULTS

### Olaparib induces immune activation in BRCA1-deficient TNBC cell lines

We compared transcriptomic analyses from four different TNBC cell lines treated with olaparib. Two cell lines were BRCA1-deficient (SUM1315 and MDA-MB-436), two were BRCA1-proficient (MDA-MB-468 and BT549), and all cell lines were *TP53*-deficient (**Fig. 1a**). Overall, BRCA1-deficient cell lines presented and shared more upregulated protein-coding genes than BRCA1-proficient cells upon treatment with olaparib (**Fig. 1b**, Additional file 1: **Fig. S1** and Additional file 2: **Table S1**). The pronounced response of BRCA1-deficient cells compared to proficient cells is consistent with *BRCA1* mutation being a predictive biomarker for olaparib therapy. Notably, shared upregulated genes in BRCA1-deficient cell lines, such as CMPK2, IFIT2, IFIT3, have been described as involved in immune response and antiviral defense (Additional file 2: **Table S2**).

**Figure 1.**
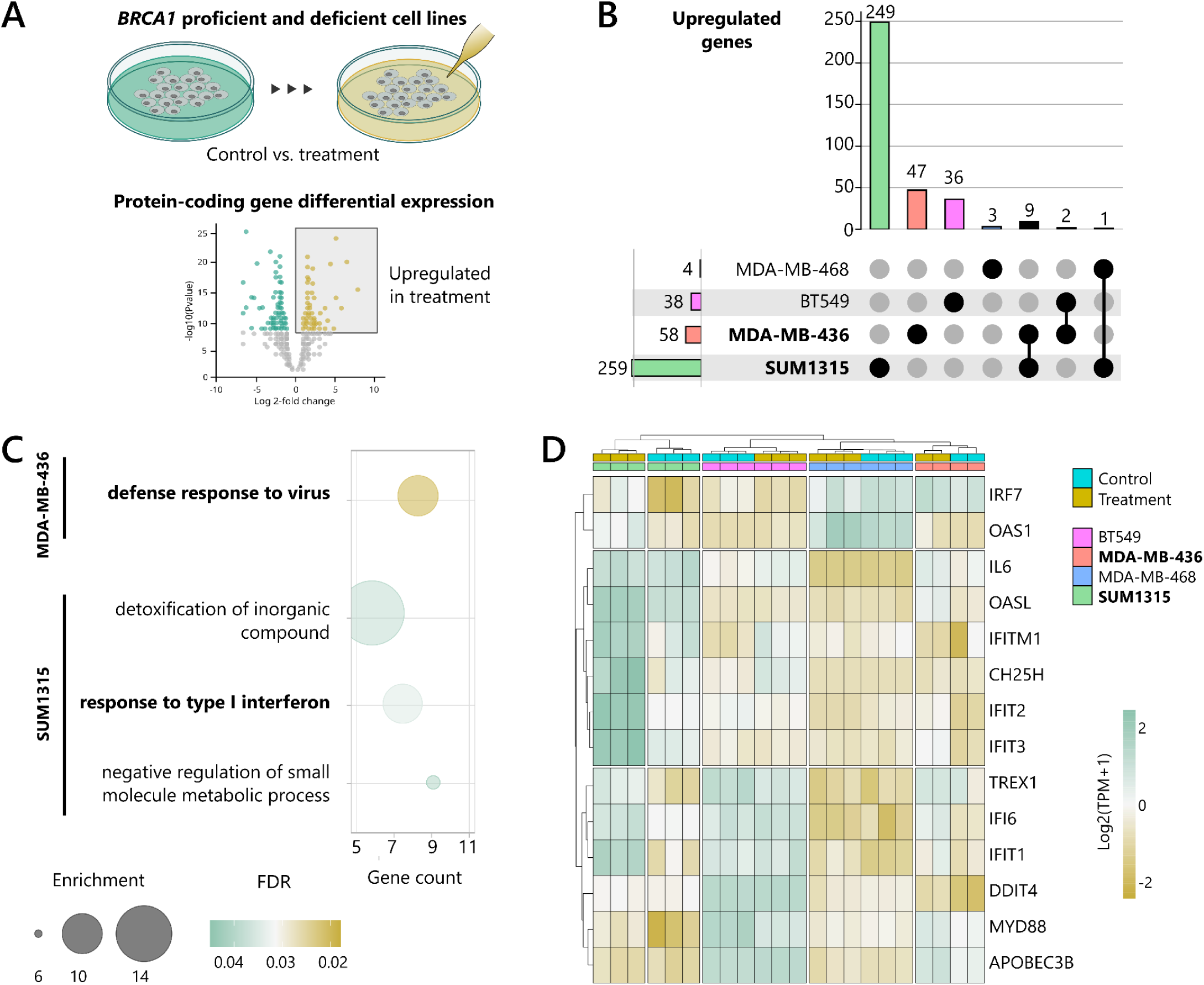
Differential analysis of protein-coding genes reveals immune activation in BRCA1-deficient cell lines upon olaparib treatment. a) Overview of the treatment and transcriptome comparison. b) Intersection of upregulated genes between cell lines (cut-off: log 2-fold change ≥ 2 and adjusted p-value < 0.05). c) Biological processes enriched in upregulated protein-coding genes in BRCA1-deficient cell lines and only displayed those with a fold enrichment higher than 6. d) Expression of genes related to immune activation across cell lines (expression values were plotted in log2 (TPM+1)). BRCA1-deficient cell lines are in bold.

Consistently, Gene Ontology (Biological Process) analysis of differentially upregulated protein-coding genes upon olaparib treatment revealed an enrichment of genes related to “defense response to virus”—a general term that includes the also enriched “response to type I interferon” process—for both BRCA1-deficient cell lines. This result was specific to the BRCA1-deficient cells and not seen for BRCA1-proficient cells (**Fig. 1c** and Additional file 2: **Table S3**). This result led us to evaluate expression changes in the immune system genes implicated in the antiviral response across all cell lines. We observed a considerable difference between treated and control samples as well as between cell lines, which were grouped accordingly (**Fig. 1d**). In both BRCA1-deficient cell lines, samples treated with olaparib showed higher expression of immune activation-related genes, including the cytosolic exonuclease TREX1 known to degrade cytosolic nucleic acids [24].

In addition, we observed a trend toward increased expression of genes within the DNA sensing cGAS-STING pathway (Additional file 1: **Fig. S2**), which can induce a type I interferon response. This trend was noted explicitly in BRCA1-deficient cell lines treated with olaparib, whereas the opposite trend was observed for BRCA1*-*proficient cells. Previous studies have demonstrated that PARP inhibition can activate the cGAS-STING pathway, thereby eliciting an immune response that may be essential for therapeutic efficacy [25,26]. Lastly members of the IFIT family, which bind 5’-triphosphate RNA, and the OAS family, which respond to double-stranded RNA (dsRNA), are upregulated by olaparib treatment in BRCA1-deficient cell lines (**Fig. 1d**). These results indicate that olaparib treatment induces an innate immune response in BRCA1-deficient cell lines, but not in BRCA1-proficient cells.

### Transposable element upregulation triggers immune response in BRCA1-deficient TNBC cells

Several studies have linked the activation of TEs as a source of an innate immune response via viral mimicry [27,28]. Mechanistically, these TE-derived RNAs can form double-stranded or repetitive structures that resemble viral intermediates and are recognized by cytosolic innate immune sensors, thereby potentially triggering an interferon-mediated antiviral response [29–32]. Accordingly, we sought to address whether the activation of transposable elements may be the source of the innate immune response induced by olaparib treatment in BRCA1-deficient cells. For this, we first investigated TE expression changes following olaparib treatment in all cell lines. We used an index built from deep-coverage long-read RNA sequencing to annotate TE transcripts more accurately, named TEscape [33]. Since TEscape uses full-length transcripts, it provides more precise and complete transcription start and termination sites, as well as transcript structural variants for all genes, including transcribed TEs. TE transcripts were classified into two main categories: TE locus and TE-TE chimera (**Fig. 2a**). TE-TE chimeras were further divided into mono- and multi-exonic types. For differential TE expression analysis, we excluded TE loci within DEGs to avoid any noise of a high expression interference in our results.

**Figure 2.**
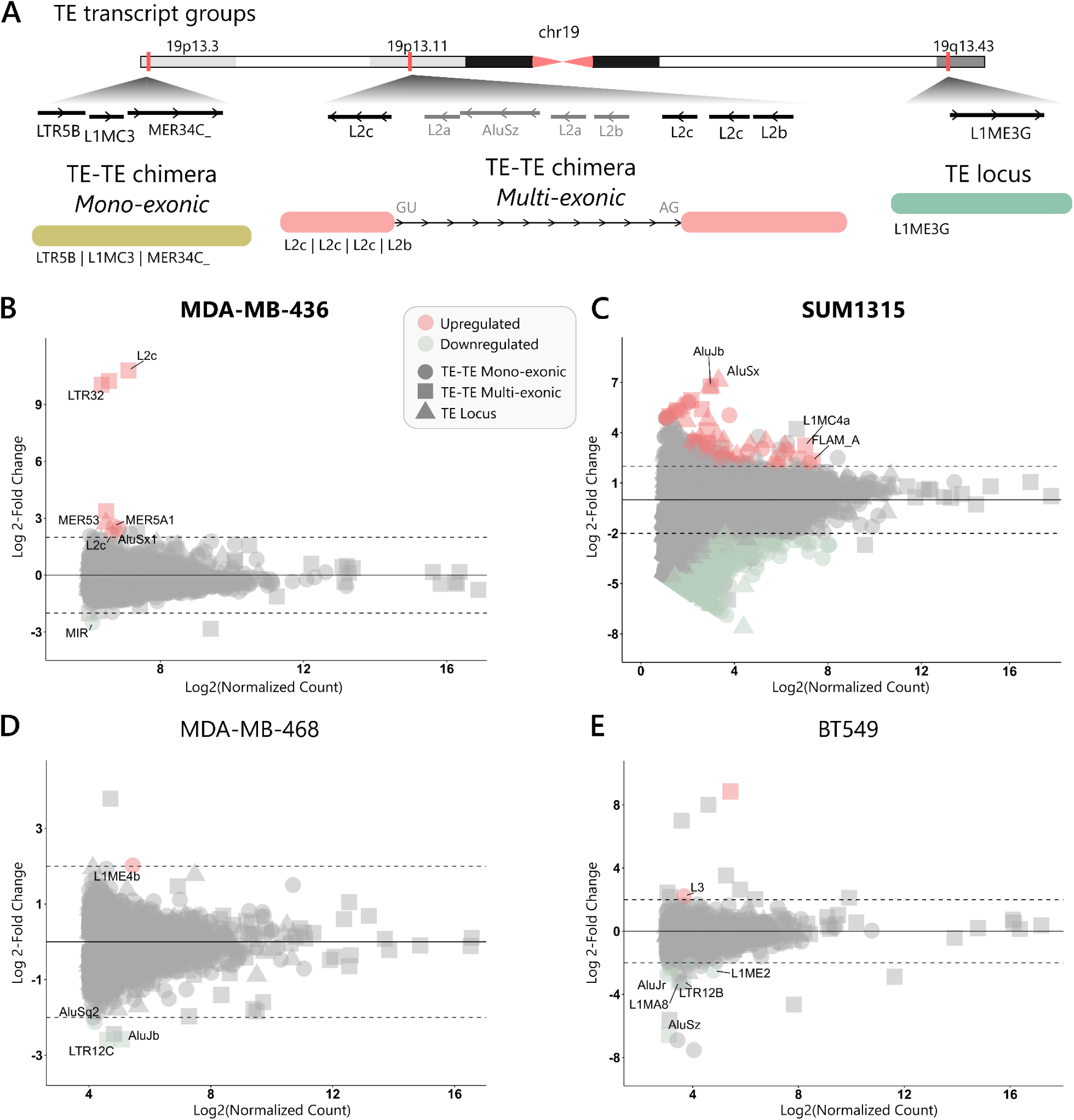
Olaparib impacts on transposable elements expression. **a)** TEscape index transcripts classification: transposable element (TE) loci and TE-TE chimeras. TE-TE chimeras are further divided into mono- and multi-exonic. The figure displays examples of transcripts present in the index. Differentially expressed TE loci and TE-TE chimeras for **b)** MDA-MB-436 (*BRCA1*^-/-^) cells, **c)** SUM1315 (*BRCA1*^-/-^) cells, **d)** MDA-MB-468 (*BRCA1*^+/+^) cells, and **e)** BT549 (*BRCA1*^+/+^) cells (cut-off: log 2-fold change ≥ 2 and adjusted p-value < 0.05). *BRCA1*-deficient cell lines are in bold.

Differential expression analysis of TEs revealed that BRCA1-deficient cell lines (**Fig. 2b-c**) presented more differentially expressed TEs transcripts, particularly, more upregulated TEs than BRCA1*-*proficient cells (**Fig. 2d-e** and Additional file 2: **Table S4**). More surprisingly, multi-exonic TE-TE chimeras were especially upregulated in BRCA1-deficient cell lines upon olaparib treatment. The top 20 upregulated TE transcripts showed a considerable difference between treatment and control samples as well as cell lines, which were grouped accordingly (Additional file 1: **Fig. S3**). These results indicate that olaparib treatment is associated with a regulation of TE expression, particularly in BRCA1-deficient cell lines.

Given that TE-derived RNAs can adopt double-stranded or repetitive structures resembling viral intermediates [29–32], we first examined the features of the upregulated multi-exonic TE-containing transcripts to address if these TE-derived RNAs can form secondary structures that can illicit an interferon response. In **Fig. 3a** we first highlighted the top 10 upregulated multi-exonic TE-TE chimera transcripts across all cell lines. Notably, the BRCA1-deficient cell line with olaparib treatment samples displayed a distinct further apart grouping from their corresponding control samples, emphasizing the impact of olaparib specifically on BRCA1-deficient cells. In both BRCA1-deficient cell lines, we observed upregulation of TE-TE chimera transcripts containing elements from the same TE subfamily, implicating the potential for the generation of intramolecular dsRNA from self-fold RNA [29–32]. Namely, distinct TE-TE chimera transcripts featuring repeats from the same subfamily—expressed in sense and antisense—point to the formation of intermolecular dsRNA from RNA duplexes [29]. These findings, coupled with the observation that olaparib treatment induced an immune response, prompted us to investigate whether the upregulated TE-TE chimera transcripts can form secondary dsRNA structures via intra- and intermolecular interactions. For this analysis, we considered mono- and multi-exonic transcripts of 200-2000 nt, as this size range aligns with the usual sizes of immunostimulatory dsRNA and TE transcripts [34]. Additional RNA folding support and cut-offs (described in the **Methods** section) were employed to confirm biological relevance and RNA stability.

**Figure 3.**
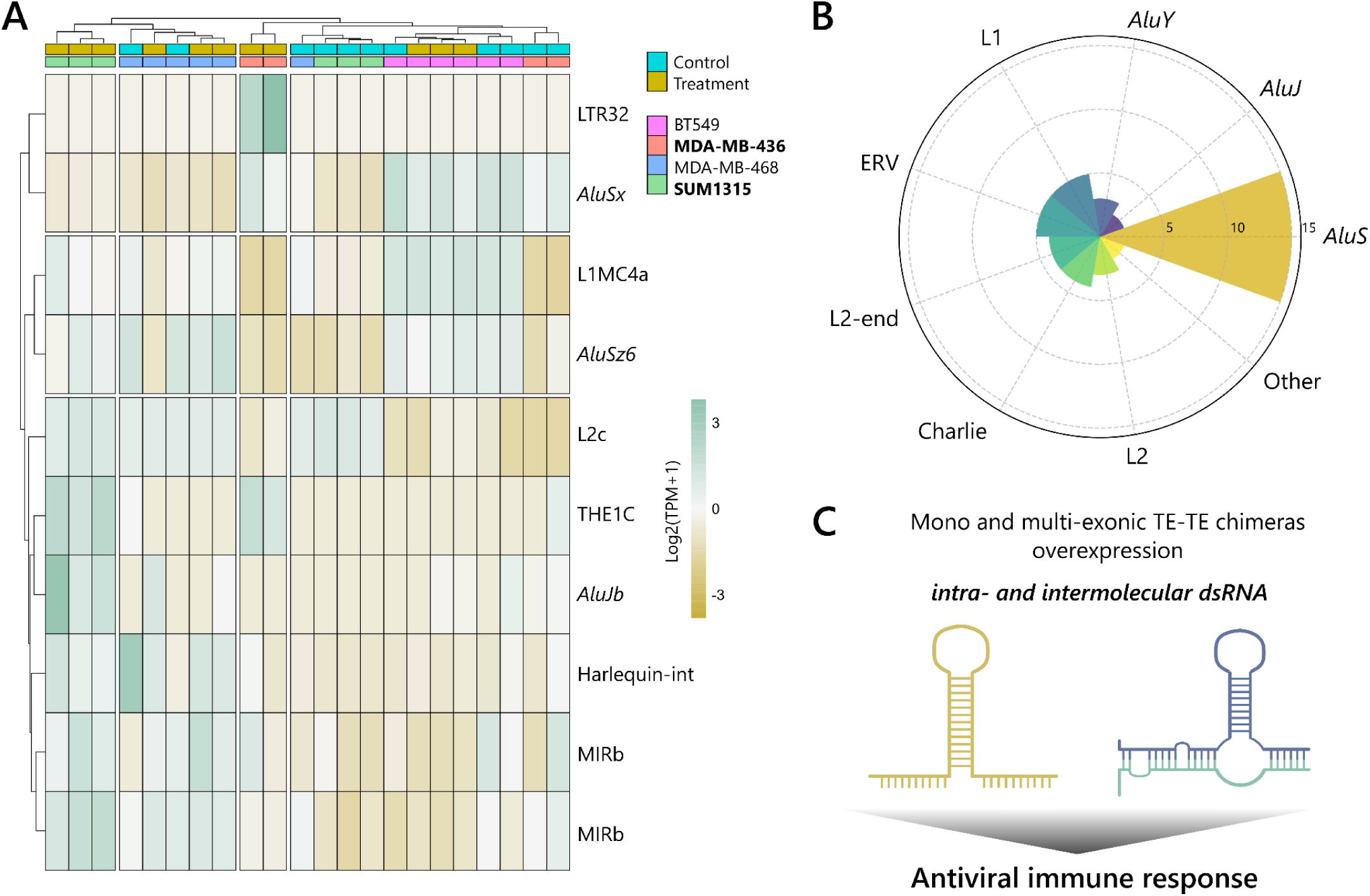
Transposable elements upregulation triggers immune response in BRCA1-deficient cells. **a)** Top 10 upregulated multi-exonic TE-TE chimera transcripts (expression values are in log2 (TPM+1)). BRCA1-deficient cell lines are in bold. **b)** Classification of TE-TE chimeras participating in intra- and intermolecular dsRNA formation. **c)** TE-TE chimeras intra- and intermolecular dsRNA formation and TE-derived RNA transcripts.

We found that both BRCA1-deficient cell lines presented TE-TE chimeras capable of forming stable intra- and intermolecular dsRNA, whereas proficient cells did not (Additional file 2: **Table S5**). Both mono- and multi-exonic TE-TE chimera transcripts were candidates of self-fold RNA, such as AluSx|AluSc8|AluSx1|7SLRNA and AluSz6|MLT1A|MER3|AluY (Additional file 1: **Fig. S4**). Mono-exonic transcripts were also capable of forming stable RNA duplexes, such as AluSx1|MER51B|AluSx3::L2c|AluSz|L2c|AluSx1 and L1M5|AluSx::L1M5|AluSx1. Consistent with previous studies [30–32], most self-folding RNA transcripts contained *Alu* repeats from the SINE subfamily (**Fig. 3b**). However, we also identified transcripts featuring repeats from other TE families, such as L1, that have not been previously reported in this context. Altogether, these findings suggest that BRCA1-deficient cell lines’ immune response to olaparib treatment might be, in part, a consequence of mono- and multi-exonic TE-TE transcripts formation of intra- and intermolecular dsRNA, in addition to other TE-derived RNA accumulating in these cells cytoplasm (**Fig. 3c**).

### Olaparib reveals transposable-element poison exons in tumorigenesis-related genes

Another known mechanism by which activation of TEs alters gene expression in various malignancies, including BC [35], is the exposure of cryptic regulatory sequences [36]. Among these newly uncovered sequences are alternative RNA splicing sites, which may generate non-coding exons. When incorporated into the mature mRNA, these exons can trigger nonsense-mediated mRNA decay; therefore, named poison exons [37]. Using the TEscape [33] index, we identified gene-TE chimeric transcripts that could represent either novel isoforms or previously annotated transcripts in Gencode [38].

Differential expression analysis of gene-TE chimeras following olaparib treatment revealed that BRCA1-deficient cell lines showed upregulation of chimeras associated with tumorigenesis. In contrast, BRCA1*-*proficient cells exhibited minimal upregulation of chimeras and lacked such function (Additional file 2: **Table S6**). The BRCA1-deficient SUM1315 cell line presented three upregulated gene-TE chimeras: *GINS1*, *GLMP*, and *ZNF169*. *GINS1* (or *PSF1*), a component of the GINS complex, is critical for DNA replication initiation and progression, playing a key role in cell cycle regulation and serving as a potential biomarker in certain cancers [39,40]. Whilst the BRCA1-deficient MDA-MB-436 cell line presented two upregulated chimeras: *BRCC3* and *ZNF10*. *BRCC3*, part of the BRCA1-A complex, is essential for the DNA damage response, particularly in the repair of DSBs [41]. For both *GINS1* and *BRCC3*, the TE segment involved in the chimeric isoforms is located in the 3’-UTR, consistent with prior isoform reports. While chimerization does not alter the protein sequence, it may introduce novel sequences that can affect gene regulation, such as microRNA binding sites [42].

Nonetheless, olaparib induced novel upregulated chimeric transcripts among Krüppel-associated box zinc finger protein (KRAB-ZNF) family members in BRCA1-deficient cells; these genes play a critical role in regulating gene expression [43]. The gene-TE chimeric transcripts arose from the exonization of poison TE exons, leading to premature termination sites (**Fig. 4a-b**). Specifically, chimerization in *ZNF169* can result in a truncated protein (**Fig. 4a**), whereas *ZNF10* chimerization cannot be translated to a protein (**Fig. 4b**). Consistently, *ZNF10* is implicated in breast invasive ductal carcinoma progression [44] and *ZNF169* promotes colorectal [45] and hepatocellular carcinoma growth [46]. Since the source genes of these gene-TE chimeras showed no differential expression in our prior analyses, we further investigated their percentage isoform composition. Chimeric isoforms of gene-TE chimeras contributed substantially to the genes’ expression in treatment samples compared to controls. While the median in control samples, the *ZNF169* and *ZNF10* chimeras accounted for 13 and 11% of isoforms, in treatment, they accounted for over 62% and 32%, respectively. Given the role of these genes in tumorigenesis, the presence of non-functional chimeric isoforms may modulate the response of BRCA1-deficient cells to olaparib therapy.

**Figure 4.**
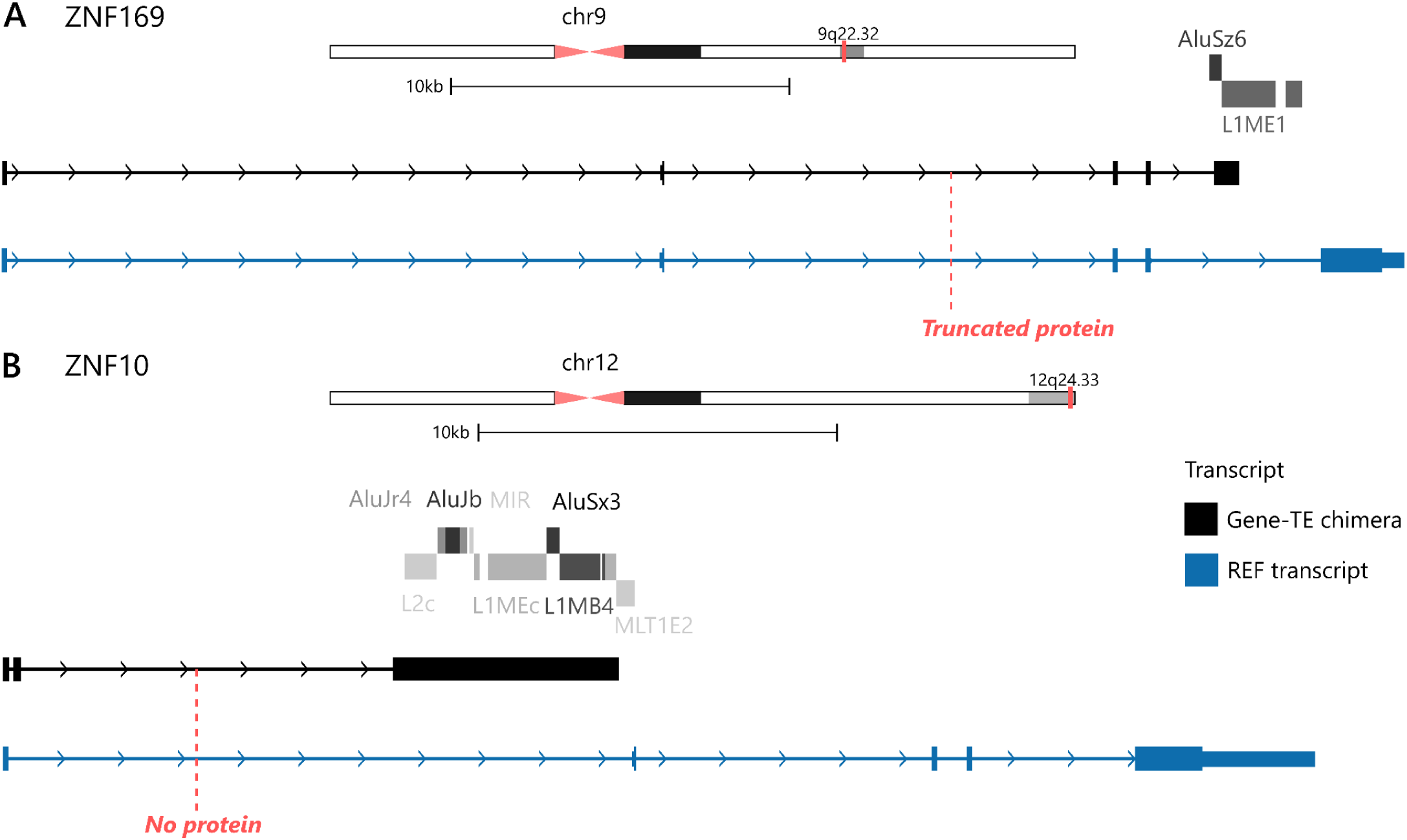
Novel gene isoforms induced by olaparib treatment uncover transposable elements’ poison exons in *BRCA1*-deficient cells. **a)** ZNF169::AluSz6|L1ME1 novel upregulated chimeric transcript in SUM1315 (*BRCA1*^-/-^) cells. Transposable element exonization results in a truncated protein. **b)** ZNF10::L2c|AluJr4|AluJb|MIR|L1MEc|L1MB4|AluSx3|L1MEc|MLT1E2 novel upregulated chimeric transcript in MDA-MB-436 (*BRCA1*^-/-^) cells. Transposable element exonization results in no protein. Both transcripts were compared to their respective reference (REF) transcript of Matched Annotation from NCBI and EMBL-EBI (MANE) [47].

### BRCA1-deficiency facilitates L1Hs activity upon olaparib treatment

Given that olaparib treatment is associated with inducing of TE expression in BRCA1-deficient cells, we hypothesized that this vulnerable genomic environment may also enable the retrotransposition of currently active mobile elements. Therefore, we performed bulk Nanopore long-read WGS in BRCA1-deficient MDA-MB-436 cells and compared control and treatment (24 h olaparib) samples. To investigate *de novo* TE insertions, we used a tool that identifies inserted segments that failed to align to the reference hg38. Since TE can also be polymorphic — common in some populations but not in others and/or absent in the reference hg38 — we used variant calls for mobile element insertions from the dbVAR database [48] to filter them out. To determine if unaligned segments indeed represented TE sequences, we performed a search against the Dfam TE sequence database [49]. We also confirmed that candidate events were absent in the control sample long-read from the same region.

We identified a 286 bp 5’-truncated L1Hs insertion at chr9 q21.11 in the olaparib-treated sample of BRCA1-deficient MDA-MB-436 cells (**Fig. 5a**). This *de novo* insertion occurred on the reverse strand, featuring hallmark characteristics of TPRT, including a poly-T tail and 5 bp target site duplications (5′-TTCTT-3′) [49]. The downstream flanking region harbored a canonical L1Hs endonuclease (EN) cutting motif on the reverse strand (5′-TTTTTA-3′) [16,50]. This insertion disrupted the intron of a novel non-coding RNA gene (ENSG00000294531) and was located just 18 bp upstream of the 3′-UTR of a long non-coding RNA (ENSG00000294571). Notably, even 5′-truncated de novo insertions — a common feature of de novo L1Hs events — can exert diverse functional consequences. These include transcript truncation [51], as observed here [52], or even the initiation of tumorigenesis [53]. This observation supports the notion that L1Hs is active in BRCA1-deficient TNBC cells following olaparib treatment.

**Figure 5.**
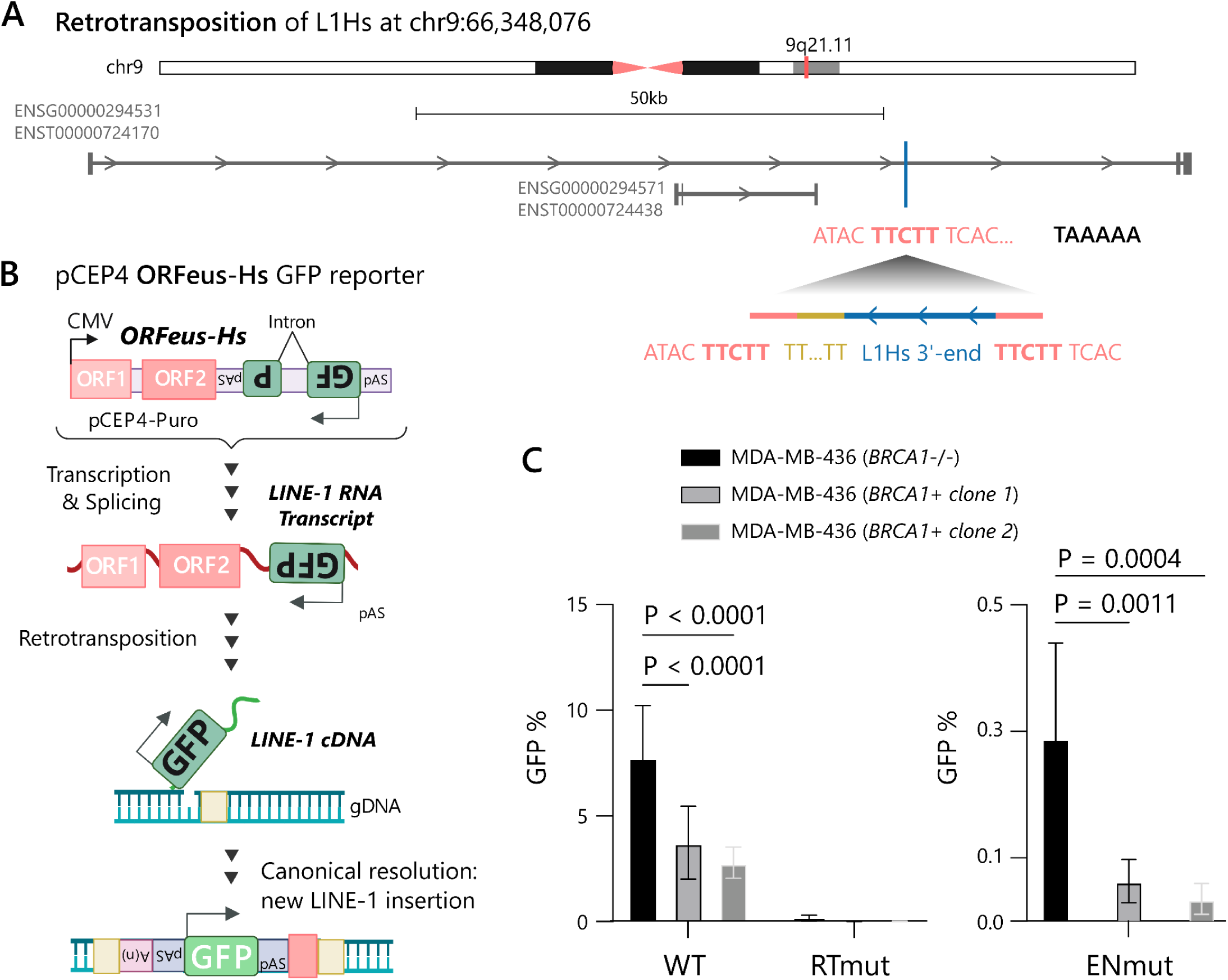
L1 activation induced by olaparib treatment in MDA-MB-436 (*BRCA1*^-/-^) cells. **a)** De novo insertion of a 5’-truncated L1Hs (blue) presenting the target site duplication (bold pink), the poly-T tail (yellow), and the endonuclease (EN) cutting motif (black). **b)** ORFeus-Hs retrotransposition GFP reporter. The ORFeus-Hs expression plasmid contains an eGFP cassette in the L1 3′-UTR which is in the opposite orientation of the L1 and is interrupted by an intron. Cells express eGFP only after the ORFeus-Hs transcript undergoes splicing and intron removal, reverse transcription, and integration into chromosomal DNA. **c)** Frequency of GFP-positive cells in MDA-MB-436 (*BRCA1*^-/-^) cells, and two MDA-MB-436 (*BRCA1*+) cell’ clones of ORFeus-Hs wild-type (WT), RTmut, or ENmut.

To directly address the impact of olaparib on L1Hs retrotransposition activity and to elucidate the role of BRCA1, we employed the synthetic L1Hs reporter construct ORFeus-Hs [54] in MDA-MB-436 cells, comparing parental BRCA1-deficient cells with cells complemented with wild-type *BRCA1* [55]. ORFeus-Hs encodes a codon-optimized human L1Hs element expressing ORF1p, ORF2p, and a GFP-based retrotransposition reporter [50,56], which we previously validated [57] (**Fig. 5b**). Unlike the WGS approach, which relies on activation of endogenous L1Hs elements, this system enables robust expression of L1Hs RNA and protein, allowing direct quantification of retrotransposition events.

We found that *BRCA1* complementation significantly suppressed L1Hs retrotransposition. The frequency of GFP-positive cells — indicative of *de novo* L1Hs insertions — was reduced by at least two-fold in the *BRCA1*-complemented cells compared to BRCA1-deficient cells (**Fig. 5c**). As expected, no GFP-positive cells were observed in cells harboring an reverse transcriptase (RT, D702Y ORF2p) deficient construct, confirming the requirement of RT for L1Hs retrotransposition as previously described [12,57,58]. Notably, we also found that *BRCA1* complementation reduced the frequency of L1Hs EN-independent insertions, which we measured using a L1Hs endonuclease (EN, H230A ORF2p) deficient reporter construct. L1Hs is known to generate de novo insertions independent of its EN activity, albeit at a relatively low efficiency compared to the EN-dependent insertions, by exploiting endogenous sites of DNA damage [16,57,59]. Together, our results show that BRCA1 suppresses both EN-dependent and EN-independent L1Hs insertions, suggesting that it likely suppresses retrotransposition independent of the first-strand nick during target-primed reverse transcription. However, it is also possible that BRCA1 deficiency increases endogenous DNA damage levels, which are then exploited for L1Hs integration independent of its EN activity. Together, our results here are consistent with several other studies showing that BRCA1 suppresses L1 retrotransposition [22,60–62].

We next sought to test the impact of olaparib treatment on the frequency of L1Hs retrotransposition in cells proficient and deficient in BRCA1. For this, we used a native L1Hs sequence (L1RP) containing a GFP-AI reporter cassette for retrotransposition in its 3’ UTR in BRCA1-proficient and -deficient RPE-1 *TP53^−^*^/−^ cells treated with increasing concentrations of olaparib. First, we found that the frequency of L1Hs retrotransposition was elevated in BRCA1*-*deficient versus BRCA1*-*proficient cells at baseline, consistent with our result above showing that BRCA1 suppresses L1Hs retrotransposition. In addition, we found that olaparib treatment suppressed L1Hs retrotransposition in a dose-dependent manner (Additional file 1: **Fig. S5**). Our result aligns with prior evidence showing that olaparib treatment inhibits EN-dependent L1Hs retrotransposition in cells [63]. Here, we found that the inhibitory effect of olaparib on L1Hs retrotransposition is independent of BRCA1, suggesting that PARP1/2 may play a critical role in resolving L1Hs insertions even in the absence of BRCA1.

### Transposable elements as hotspots of unconservative repair mechanisms

Interestingly, a duplication event was initially misclassified as a de novo L1Hs insertion, owing to the presence of an L1PA2 element detected through Nanopore long-read whole-genome sequencing in BRCA1-deficient MDA-MB-436 cells treated with olaparib. In further detail, this event was a non-tandem duplication of a pre-existing 5’-truncated L1PA2 of 559 bp at chr11 p11.12 (**Fig. 6a-b**), exclusively observed in the olaparib-treated sample and absent in the control. We used the BLAT tool [64] from UCSC Genome Browser [65] to search for sequence similarity elsewhere in the genome to the newly inserted sequence and found the same L1PA2 upstream in the chromosome (∼190 Mb distance) (**Fig. 6a**). We confirmed the sequences’ high similarity using Blast2Seq [66], which also revealed a template-switching (**Fig. 6b**). L1PA2 duplication occurred in the region of two different transcript isoforms from the same lncRNA gene (LINC02750). Looking further into this, we also found a 15 bp deletion at the duplication site, and a 14 bp microhomology between the duplication site and the original L1PA2 flanking region. The identification of a non-tandem duplication event with template switching, accompanied by a small deletion and microhomology, suggests that the duplication may be a result of a repair by microhomology-mediated break-induced replication (MMBIR) [67]. These findings shed light on the underlying repair mechanisms at play in BRCA1-deficient TNBC cells upon olaparib treatment and on TEs as hotspots for unconservative repair mechanisms.

**Figure 6.**
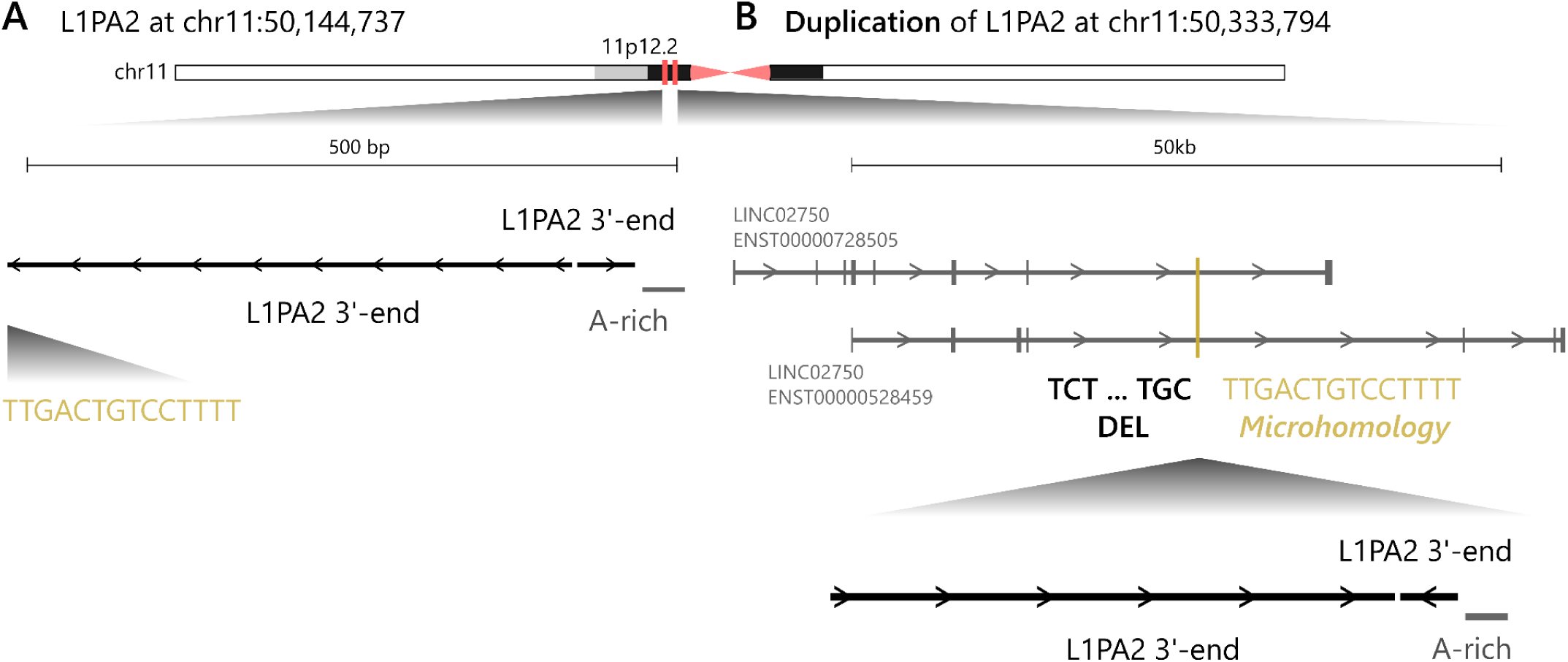
Duplication of an L1PA2 element induced by olaparib treatment in MDA-MB-436 (*BRCA1*^-/-^) cells. **a)** L1PA2 is present in the reference genome, and its **b)** Duplication is further away in the genome, presenting a deletion (black), a microhomology (yellow), and the template switching.

### Olaparib and the expression of young L1 families

Transposition cannot always be attributed to specific progenitor elements since quantifying L1 RNA at specific loci poses significant challenges [68]. When characterizing TE expression using TEscape, this limitation persisted due to the locus-specific methodology applied. Nevertheless, identifying the expression of L1 loci is crucial for understanding the basis of retrotransposition upon olaparib treatment. To address these challenges, we employed a complementary approach using L1EM [69], which enables loci-specific quantification of L1 transcripts. L1EM categorizes L1 transcripts into four distinct groups and also separates transcripts that are capable of generating retrotransposition from those that are not (see **Methods** for further details).

To deepen our understanding of how olaparib could stimulate L1 activity, we investigated L1 locus-specific expression in BRCA1-deficient MDA-MB-436 cells (as in our previous results). Our analysis focused on young L1 families — specifically L1Hs and L1PA2 — because these families are known to harbor active, autonomous loci [13,69]. We identified deregulation of a few L1PA2 and L1Hs loci, highlighting how challenging it is to recover L1 loci-specific expression (Additional file 1: **Fig. S6** and Additional file 2: **Table S7**). Although our analysis showed a few down-regulated L1 loci, they are, in fact, passive transcripts not driven by L1 expression. Still, we found an interesting full-length (6026 bp) L1PA2 candidate. Overall, these results indicate that olaparib influences the expression of young L1 families, thereby establishing a foundation for our findings on L1 activation upon olaparib treatment.

## DISCUSSION

Here, we presented a comprehensive characterization and investigation of BRCA1-deficient and BRCA1-proficient TNBC cells’ response to olaparib. Our findings reveal a BRCA1-dependent immune activation: olaparib treatment selectively induces an antiviral-like response in BRCA1-deficient TNBC cells, marked by the upregulation of genes involved in the detection and processing of RNA species—including dsRNA, ssRNA—as well as dsDNA and ssDNA. This immune response appears to be driven by the overexpression and cytoplasmic accumulation of TE-derived transcripts BRCA1-deficiency creates a permissive environment for TE expression and L1Hs retrotransposition, which, in turn, may drive immunostimulatory signaling and DNA damage. Concurrently, we identify additional genomic vulnerabilities in BRCA1-deficient TNBC cells, where structural variation is tightly linked to the repetitive nature of TEs, suggesting that the sequences of these elements serve as hotspots for non-conservative DNA repair and further genomic instability.

Our results demonstrate that BRCA1 suppresses L1Hs retrotransposition in TNBC cells. Namely, BRCA1 deficiency causes an elevation of both EN-dependent and -independent L1 insertions. Mechanistically, our results suggest that BRCA1 suppresses L1 retrotransposition downstream of the first-strand DNA nick during target-primed reverse transcription. Our results are consistent with several genetic screen studies showing a critical role of BRCA1 in suppressing L1 retrotransposition [22,60,70,71], which have proposed a working model of BRCA1 in preventing L1 integration at ongoing DNA replication forks. We additionally suggest here that the complex DNA lesions caused directly or indirectly by olaparib treatment might serve as vulnerable sites of L1 insertions. Here, we found that olaparib inhibits EN-dependent L1 insertions in both BRCA1-proficient and BRCA1-deficient cells, consistent with a previous study showing that PARP1/2 are required for L1 retrotransposition [63]. However, olaparib suppressive effect on L1 retrotransposition appears specific on EN-dependent insertions, but not EN-independent insertions [63]. In view of this, we propose that olaparib-induced unrepaired DNA lesions can be resolved through retrotransposon-mediated DNA repair, thereby bypassing the requirement for EN and PARP2 for retrotransposition (**Fig. 7**). Additionally, BRCA1 may regulate L1Hs expression via interactions with the SWI/SNF chromatin remodeling complex [72] and the DNA methyltransferase DNMT1 [21]. Collectively, our findings indicate that BRCA1-deficient cancer cells can sustain L1Hs expression and retrotransposition following olaparib treatment, in stark contrast to its dose-dependent suppressive effects on L1Hs activity in normal cells.

**Figure 7.**
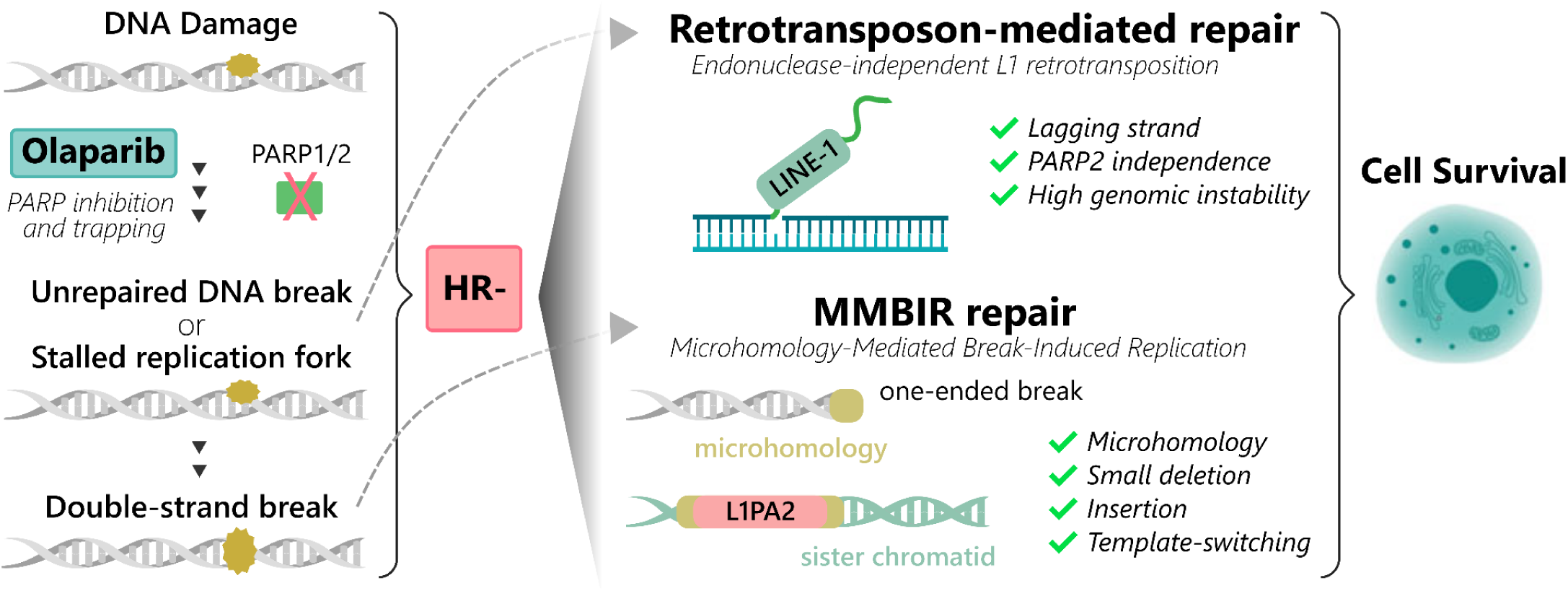
Repair mechanisms induced by olaparib in BRCA1-deficient cancer cells. Olaparib-induced PARP inhibition and trapping lead to stalled replication forks (S-phase) and unrepaired DNA breaks. Lagging strands generated in this context may serve as primers for LINE-1 reverse transcriptase, facilitating retrotransposon-mediated repair through EN-independent retrotransposition. Concurrently, replication fork collapse and unrepaired DNA damage can produce double-strand breaks (DSBs), particularly one-ended DSBs in highly unstable genomic environments. During S-phase, these breaks may be repaired via microhomology-mediated break-induced replication (MMBIR), an error-prone mechanism that promotes structural variation. Together, retrotransposon-mediated repair and MMBIR contribute to cancer cell survival under therapeutic pressure.

Simultaneously, olaparib can cause complex DNA lesions including PARP trapping on DNA, olaparib-induced replication fork collapse and unrepaired single-strand breaks, all of which can result in toxic DSBs [23,67]. The generation of free DNA ends occurs, for instance, in regions where there is no replication fork coming from the opposite direction. Given that PARPi-induced collapsed forks such as single-ended DNA breaks cannot be repaired by non-homologous end joining (NHEJ) [73], we propose that MMBIR is engaged to resolve these breaks. When equal loci on sister chromatids or homologous chromosomes are available (S-phase) and selected as a donor, the resolution of the DNA break is neutral. Conversely, when HR repair is compromised, MMBIR may be activated in response, utilizing short homologous sequences to resolve these breaks (**Fig. 7**). This repair facilitates misalignment of non-allelic loci. The involvement of an L1PA2 indicates that repetitive elements may be particularly susceptible to non-allelic homologous alignment [67], thereby promoting structural variations [74] including duplications and microdeletions.

Interestingly, our findings align with the dual role of TE in cancer [75]. While TEs may act as hotspots of PARPi resistance, they also expose genomic vulnerabilities that can be therapeutically exploited under the same conditions. Encouraging findings about TEs deregulation in response to therapies for which they are not the primary target have sparked scientific interest [75,76], including in TNBC [77]. Further research could enlighten if the olaparib-driven immune response in BRCA1-deficient cells can sensitize tumors to immunotherapies, similarly to other TE-activating drugs [78–80]. Taken together, our findings provide new insights into olaparib’s synthetic lethality and previously unrecognized mechanisms of TEs.

## METHODS

### Reagents and Cell Culture

Olaparib (AZD2281) was purchased from Selleckchem (Houston, Texas, USA) and was dissolved in dimethyl sulphoxide (DMSO). BRCA1-deficient MDA-MB-436 TNBC cells were obtained from Hospital Sírio-Libanês (São Paulo, Brazil), cell line identity was confirmed by short tandem repeat analysis (STR) and absence of mycoplasma contaminations by PCR. Cells were maintained in RPMI supplemented with 10% fetal bovine serum and used within 3-8 passages. Cells were kept at 37°C, with a minimum relative humidity of 95% and an atmosphere containing 5% CO_2_. MDA-MD-436 cells engineered to express BRCA1-proficient [55], as well as control cells containing an empty vector, were a generous gift from Dr. Bing Xia (Rutgers Cancer Institute). RPE-1 *TP53*^-/-^ hTERT and RPE-1 *TP53*^-/-^ *BRCA1*^-/-^ hTERT [81] were a generous gift from Dr. Dipanjan Chowdhyury (Dana-Farber Cancer Institute).

### Cell Survival

MDA-MB-436 cells were seeded at 1 - 1,5 × 10³ cells/well in 96-well plates and treated with olaparib at 0-64μM concentrations. Cell viability was assessed at 72h post-treatment by the Resazurin assay. Data were expressed as a percentage of control and were the mean±SD of four separate experiments, each performed in triplicate. Half maximal inhibitory concentration (IC50) of olaparib was determined by variable slope least squares fit of inhibitor vs. normalized response as 11.72µM (rounded up to 12µM), and this concentration was used for the experiments (Additional file 1: **Fig. S7**).

### RNA extraction and sequencing

Cells were seeded at 40000 cells per well in 6-well plates, after 24h of acclimation, submitted to 24h of control/treatment. Control samples were incubated for 24h in DMSO, and treatment samples for 24h in 12uM olaparib. RNA sequencing (RNA-seq) was performed using poly-A + NEBNext Ultra II Directional Library in a DNBSEQ-T7 PE150 machine. The RNA-seq data were deposited in the European Nucleotide Archive (ENA)(PRJEB97215).

### RNA-seq data processing

We used RNA-seq data from the BRCA1*-*proficient MDA-MB-468 and BT549 TNBC cell lines and the BRCA1-deficient TNBC SUM1315, available in GenBank (NCBI) under accession GSE165914 and GSE149621. All samples were prepared as single-end triplicate libraries. MDA-MB-436 samples were in duplicate paired-end libraries. Comparisons were between treatment and control samples for each cell line. The treatment consisted of the IC50 of each cell, specifically, overnight exposure to 3µM olaparib for MDA-MB-468 and 8µM for SUM1315, 48h of exposure to 20µM olaparib for BT549, and 24h exposure to 12µM olaparib for MDA-MB-436. Data were downloaded from the Sequence Read Archive using the SRA toolkit v3.0.0 (https://github.com/ncbi/sra-tools) and converted to FastQ with fastq-dump. FastQC v0.11.9 software [82] was used to assess the sequencing quality of the data. Reads were trimmed using a custom Pearl script if necessary.

### Gene and TE expression

TEscape v1.0 [33] index was merged with Gencode v36 [38] genes. Kallisto v0.48.0 [83] was used for pseudo-alignment and transcript-level read quantification. Gene transcripts were aggregated to the gene level using the tximport v1.3.9 [84] R package. The merged gene and TEs count tables were filtered to include only samples with either all control or all treatment samples differing from zero, and submitted to DESeq2 v1.38.3 [85]. Differentially expressed TE loci within DEGs were dropped. Cut-off of log 2-fold change ≥ |2| and adjusted P-value of 0.05 were used to consider genes, TE transcripts, and gene-TE chimeric transcripts as differentially expressed. For better visualization purposes, TE-TE chimera names were displayed based on the first TE participating in the transcript (supplementary tables display the name of all participants). The Gene Ontology Resource v20240807 [86] was used for enrichment analysis of DEGs (PANTHER Overrepresentation Test, Fisher test type, and FDR correction). Biological processes were considered enriched if fold enrichment was over 2 and FDR < 0.05. Gene-TE chimeric transcript putative protein formation and comparison with the canonical transcript protein ORF finder [87] from NCBI was used.

### TE-TE chimera transcripts dsRNA formation

We employed the ViennaRNA [88] package for the analyses of intra- and intermolecular dsRNA formation. RNAfold v2.6.3 was used for intramolecular RNA folding, and RNAcofold v2.6.3 was used for intermolecular RNA duplexes. Cut-offs included: upregulated TE-TE chimeras with transcript sizes varying from 200 to 2000 nt and ΔG/nt < -0.25 kcal/mol. In addition, for intramolecular dsRNA, a Z-score compared to that of 100 randomized sequences ((energy - mean energy) / standard deviation energy < -3) and, for intermolecular RNAs, the ensemble ΔG of the heterodimer should be lower than that of the homodimers.

### DNA extraction and sequencing

Cells were seeded at 200 x10^6^ cells per bottle, after 24h of acclimation, and submitted to 24h of control/treatment. The Ultra-Long DNA Sequencing Kit V14 (SQK-ULK114) was used in combination with the Monarch HMW DNA Extraction Kit for Tissue (New England Biolabs, T3060) to extract ultra-high molecular weight gDNA. Sequencing was performed in a PromethION (Oxford Nanopore Technologies) machine. The median sequencing depth across chromosomes (1-22) was 16.68x for the control sample and 14.53x for the treatment sample, with a median percentage of covered bases of 95% in both samples. The median sequencing depth of coverage per chromosome is displayed in Additional file 1: **Fig. S8**. The WGS data were deposited in ENA (PRJEB97215).

### Whole-genome insertions calling

NanoPlot and samtools coverage were used to assess read quality. Reads filtered for “PASS” were mapped to the reference genome hg38 with minimap2 v2.24-r1122. BAM files of each fastQ were merged with Samtools merge. Samtools MAPQ was used to keep reads with a Phred Quality Score of 20 or higher. The GraffiTE tool [89] was used to identify unaligned segments to the reference genome, and the Dfam TE sequence database to identify these segments as TEs. For this analysis, polymorphic insertions were filtered using variant calls of mobile element insertions from the dbVAR database [48]. The absence of de novo insertion candidates in the control sample was confirmed, and all candidates identified in the treatment sample were manually validated.

### L1Hs retrotransposition reporter assays

#### MDA-MD-436 cells

MDA-MD-436 cells complemented with BRCA1-proficient (two clones) and the control cells containing an empty vector were established and validated by Dr. Bing Xia [55]. To assay L1Hs retrotransposition frequency, we used a synthetic L1Hs sequence (ORFeus-Hs wild-type) [54] encoding ORF1p and ORF2p and containing a GFP reporter for retrotransposition in the 3’ UTR (pMT527). We also used two retrotransposition mutant versions of the L1Hs reporter plasmid, each containing mutations in the encoded ORF2p: a reverse transcriptase mutant (pLD631, D702Y) and an endonuclease mutant (pLD632, H230A). These L1Hs constructs were expressed from a pCEP4 episomal vector modified to contain a Puromycin resistance gene under a CMV promoter. All three plasmids have been previously validated [57].

We used these plasmids to perform the L1Hs GFP-AI reporter assay for retrotransposition in the *BRCA1* complemented and control MDA-MD-436 cells as previously described [57,90]. Briefly, we seeded 1 x10^5^ cells in 6-well plates (day 1, d 1). The following day, we transfected each well with Viafect (Promega) and 1 µM of each pCEP4-Puromycin plasmid (d 2): pMT527 (wild-type), pLD631 (RTmutant-D702Y), pLD632A (ENmutant-H230A). The media for cells transfected with the L1Hs reporter plasmids was changed again two days after transfection (d 4) and supplemented with 5µM/mL of Puromycin. Cells were then incubated until day 7 when cells were collected and assayed for the percentage of GFP+ cells by flow cytometry (BD LSRFortessa). In addition, a pCEP4-Puromycin vector expressing a GFP cassette (MT498, pCEP4-GFP) was used to monitor transfection efficiency by assaying the percentage of GFP+ cells on d 4 by flow cytometry (BD LSRFortessa). We normalized the percentage of GFP+ cells from the cells transfected with the L1Hs reporter plasmids to the percentage of GFP+ cells from cells transfected with the GFP plasmid.

#### RPE-1 hTERT cells

We also performed the L1Hs-GFP reporter assay in RPE-1 *TP53*^-/-^ hTERT and in RPE-1 *TP53*^-/-^ *BRCA1*^-/-^ cells, which were previously established by Dr. Dipanjan Chowdhury [81]. For this experiment, we used a native L1Hs sequence (L1RP, GenBank: AF148856.1) with its 3’UTR modified to contain a GFP-AI reporter cassette for retrotransposition, which we previously validated [90] and was expressed from a pCEP4 vector modified to express Puromycin instead of Hygromycin. The L1Hs-GFP reporter assay as above, except the media containing olaparib or DMSO as a control was supplemented at day 4.

#### Young L1 elements expression

L1EM v1.1 [68] was used to quantify L1Hs and L1PA2 at the genomic locus level. L1EM uses an expectation-maximization algorithm to separate transcripts capable of generating retrotransposition from those that are not. Therefore, L1EM categorizes L1 transcripts into four distinct groups: (1) ‘Only’ transcripts: these run sense and include only the annotated L1 element. (2) ‘Run-on’ transcripts: these run sense and extend into downstream sequences. (3) ‘Passive’ transcripts: these encompass both upstream and downstream sequences and are further divided into sense and antisense categories. (4) Antisense transcripts: these run antisense and include the first 500 bases of the L1 element, along with upstream sequences. The count table was filtered to include only samples with either all control or all treatment counts different from zero, and submitted to DESeq2. Cut-off of log 2-fold change ≥ |2| and adjusted P-value of 0.05 were used to consider L1 elements as differentially expressed.

## SUPPLEMENTARY INFORMATION

Additional file 1: PDF document containing Figures S1–S8.

Additional file 2: Microsoft Excel document containing Tables S1–S7.

## LIST OF ABBREVIATIONS

BC: Breast Cancer
TNBC: Triple-Negative Breast Cancer
HR: Homologous Recombination
TE: Transposable Element
L1Hs: Long Interspersed Nuclear Element 1 human-specific
PARPi: PARP inhibitor
DSB: Double-Strand DNA Break
WGS: Whole-Genome Sequencing
DEG: Differentially Expressed Gene
dsRNA: double-stranded RNA
dsDNA: double-stranded DNA
ssDNA: single-stranded DNA
EN: Endonuclease
RT: Reverse Transcriptase
MMBIR: Microhomology-Mediated Break-Induced Replication
NHEJ: Non-Homologous End Joining
RNA-seq: RNA sequencing

## DECLARATIONS

### DATA AVAILABILITY

The datasets supporting the conclusions of this article are available in the European Nucleotide Archive (ENA) repository, under BioProject PRJEB97215 (https://www.ebi.ac.uk/ena/browser/view/PRJEB97215), and in GenBank (NCBI) under BioProjects GSE165914 (https://www.ncbi.nlm.nih.gov/bioproject/?term=GSE165914%20) and GSE149621 (https://www.ncbi.nlm.nih.gov/bioproject/?term=GSE149621%20).

### AUTHOR’S CONTRIBUTIONS

D.M.M., J.C.O., E.L.S.L. and P.A.F.G. conceived the project. D.M.M., J.C.O. and E.L.S.L. designed cell line experiments, transcriptome and whole-genome sequencing. D.M.M. performed cell line experiments for transcriptome sequencing and analyses of experimental data with support from S.C.S.B., J.C.O. and E.L.S.L. D.M.M. and M.A.P. performed cell line experiments for whole-genome sequencing with support from J.C.O. and E.L.S.L. M.A.P. performed whole-genome sequencing with support from J.C.O. C.M.-D. and P.S. performed ORFeus-Hs experiments with support from K.H.B. D.M.M. performed all the bioinformatic analyses with support from R.L.V.M and P.A.F.G. D.M.M. and P.A.F.G. wrote the manuscript with input from all authors.

