## Additional File 1 for "Transposable elements shape olaparib response according to BRCA1 status in triple-negative breast cancer"

### **Additional File 1:** Supplementary Figures S1-S8

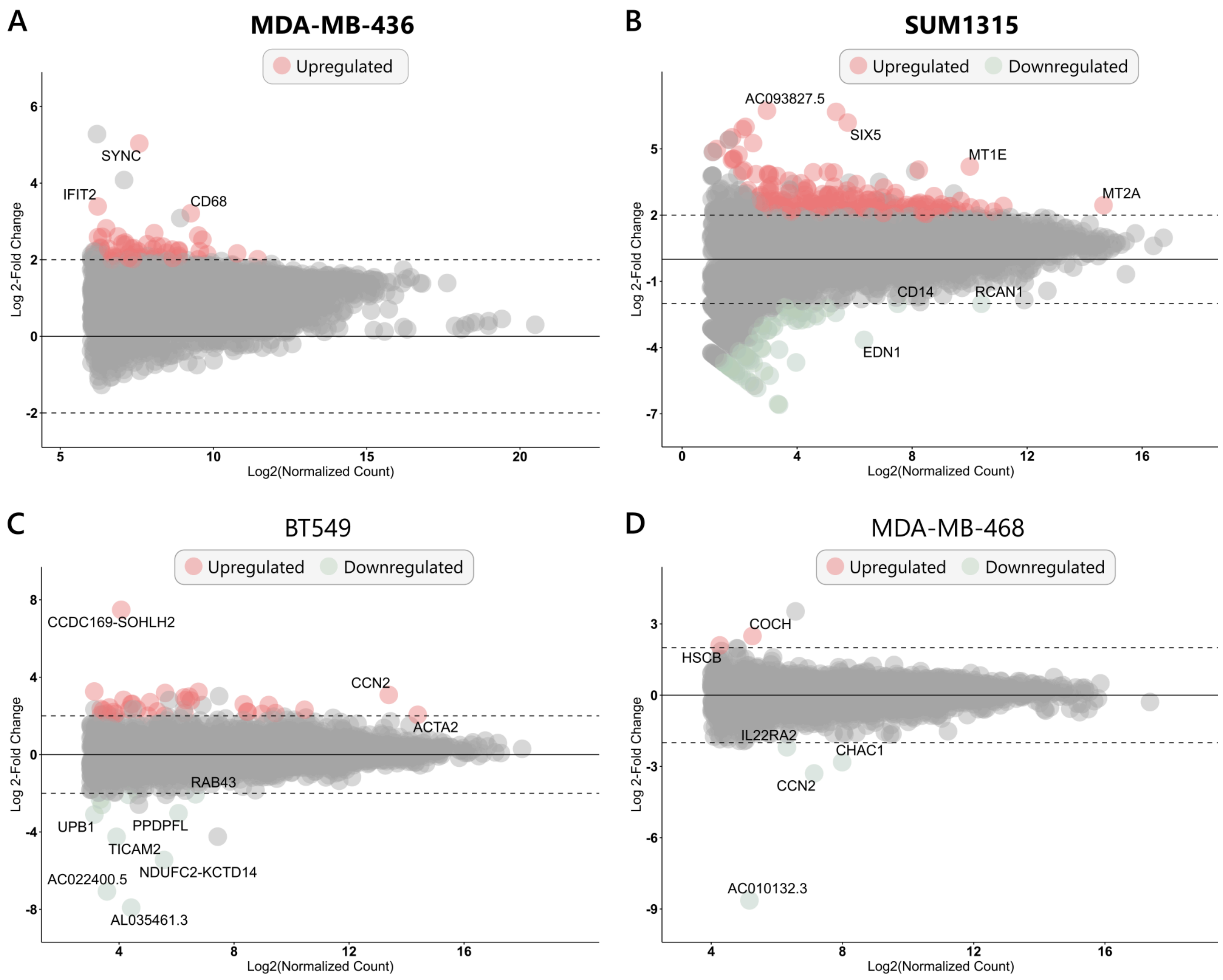


**Supplementary Figure S1. Differential expression of protein-coding genes upon olaparib treatment for a)** MDA-MB-436 (BRCA1$^{-/-}$) cells, **b)** SUM1315 (BRCA1$^{-/-}$) cells, **c)** MDA-MB-468 (BRCA1$^{+/+}$) cells, and **d)** BT549 (BRCA1$^{+/+}$) cells (cut-off: log 2-fold change ≥ 2 and adjusted p-value < 0.05). *BRCA1*-deficient cell lines are in bold.

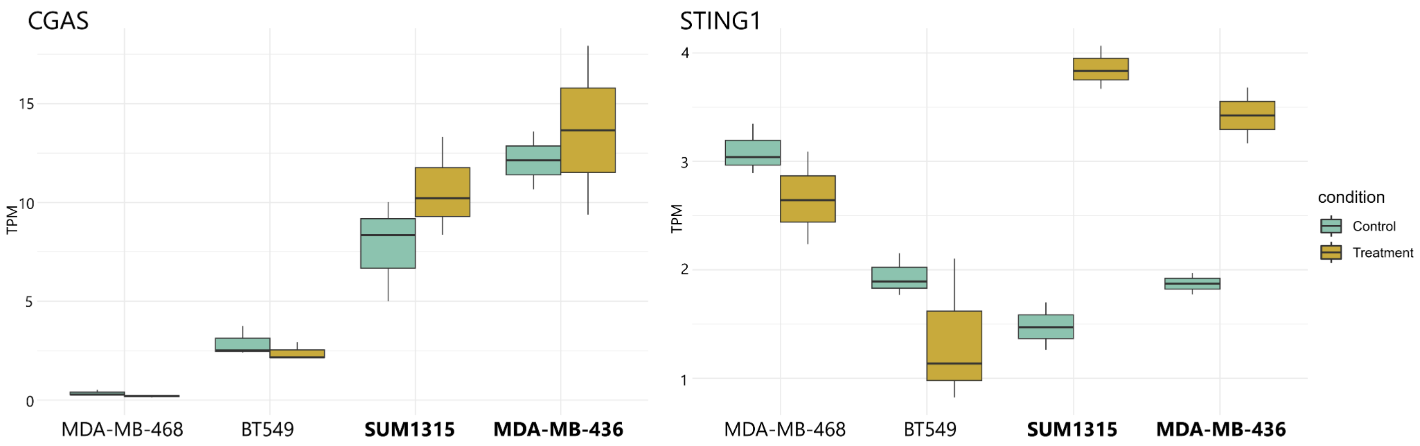


**Supplementary Figure S2. Expression in TPM of *CGAS* and *STING1* in control (light green) and treatment (olaparib; mustard) samples across cell lines.** *BRCA1*-deficient cell lines are in bold.

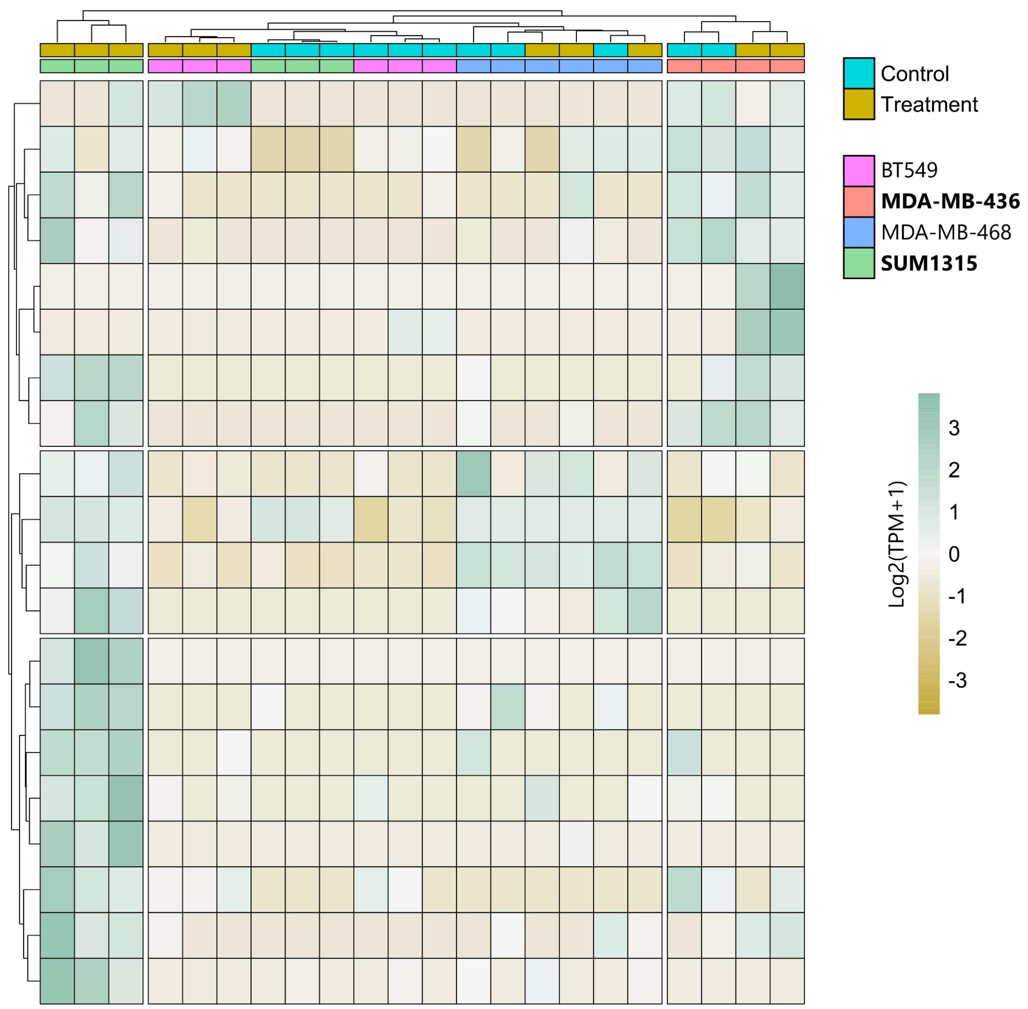


**Supplementary Figure S3. Top 20 upregulated transposable elements transcripts.** Expression values are in Log2 (TPM+1). *BRCA1*-deficient cell lines are in bold.

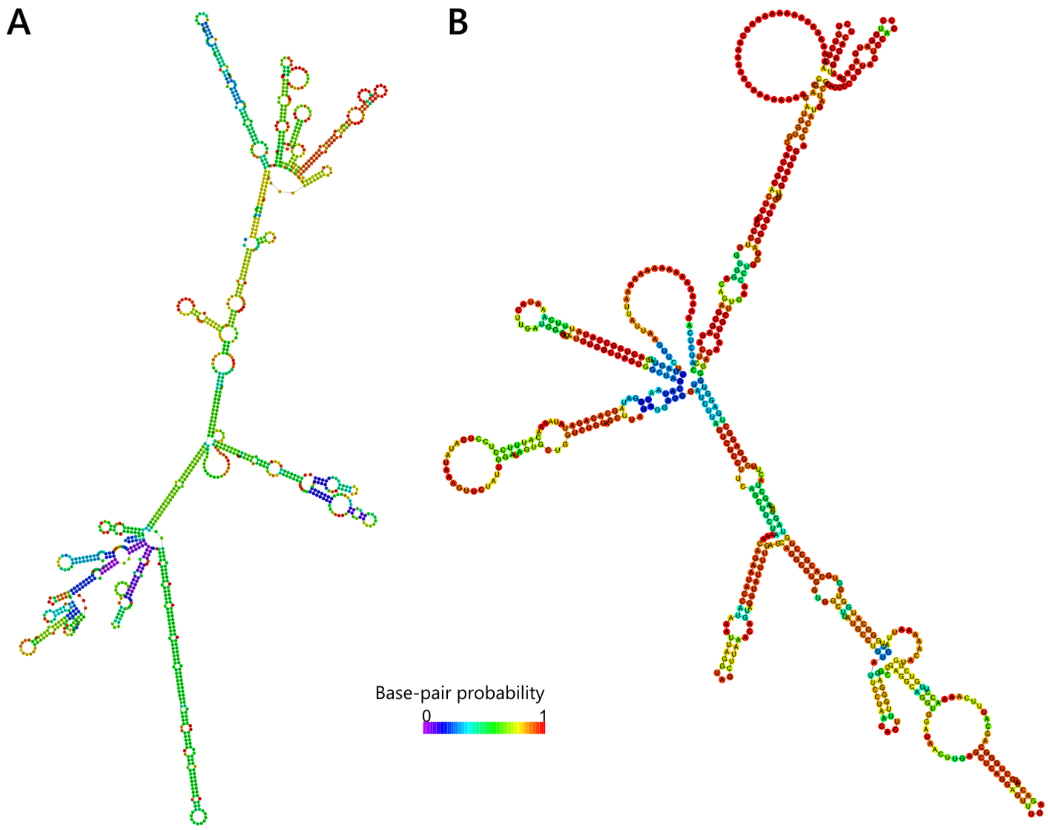


**Supplementary Figure S4. Intramolecular double-stranded RNA formation by TE-TE chimera transcripts. a)** Multi-exonic TE-TE chimera AluSx|AluSc8|AluSx1|7SLRNA displaying self-fold dsRNA. **b)** Mono-exonic TE-TE chimera AluSz6|MLT1A|MER3|AluY displaying self-fold dsRNA. Color coded by base-pair probability.

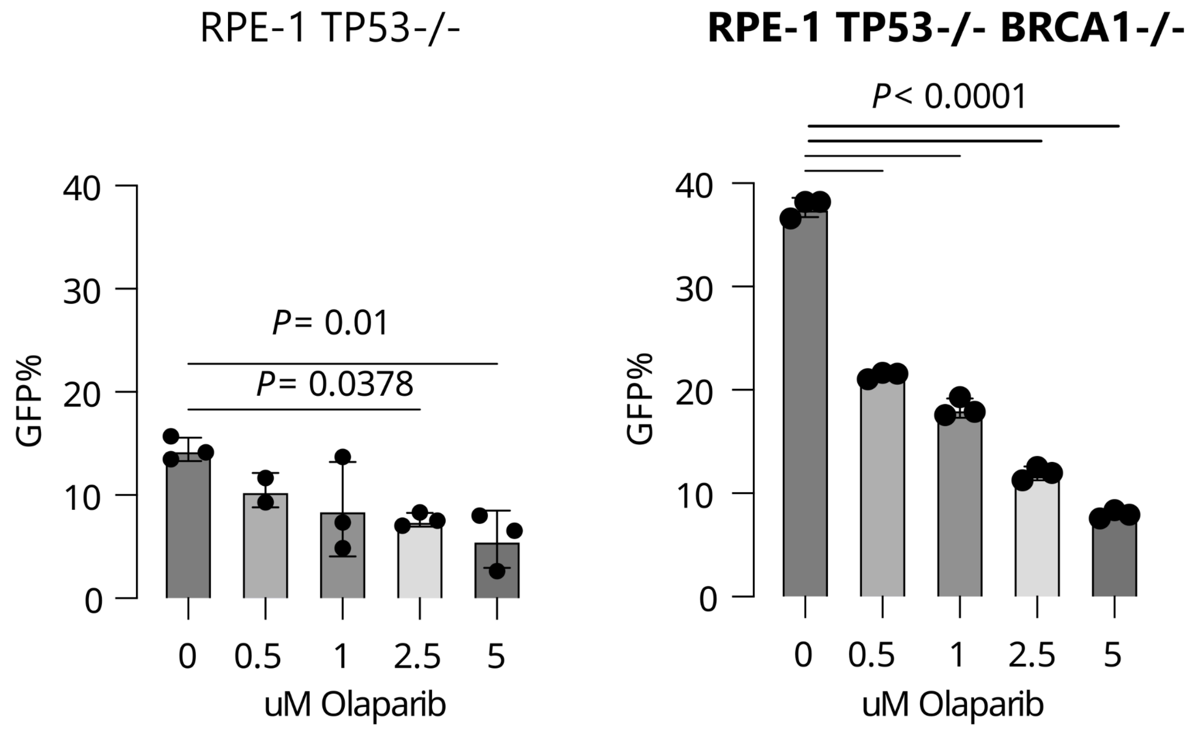


**Supplementary Figure S5. Frequency of GFP expressing cells in RPE-1 (*BRCA1*$^{+/+}$) cells, and RPE-1 *BRCA1*$^{-/-}$ (*BRCA1*-depleted) cells of ORFeus-Hs.**

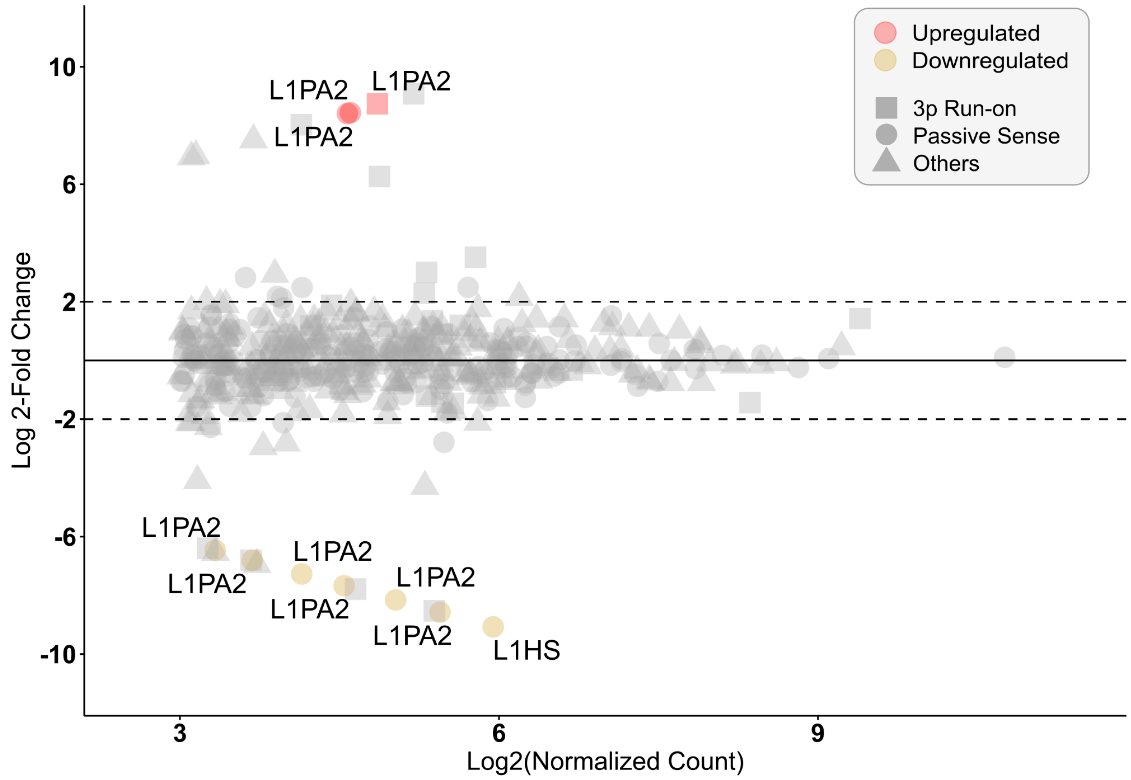


**Supplementary Figure S6. Young L1 loci-specific differential expression under olaparib treatment in MDA-MB-436 (BRCA1$^{-/-}$) cells.** Cut-off: log 2-fold change ≥ 2 and adjusted p-value < 0.05.

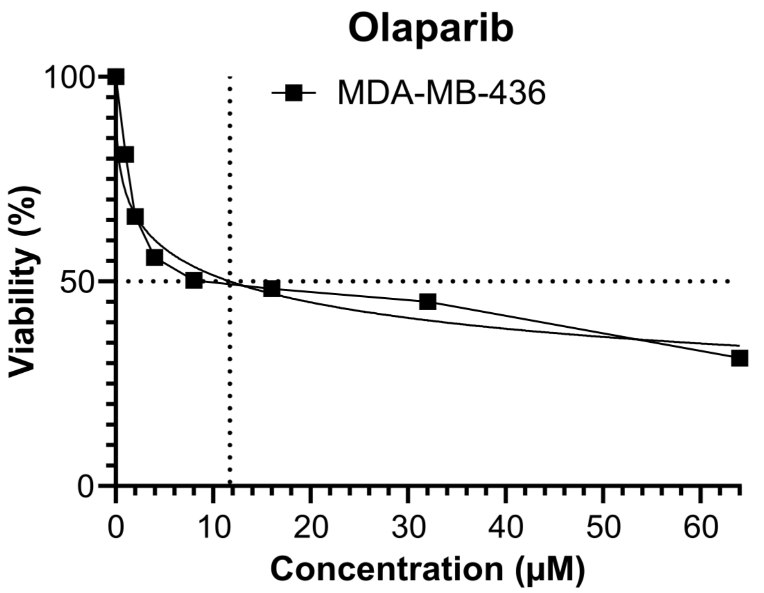


**Supplementary Figure S7. Half maximal inhibitory concentration (IC50) of olaparib in MDA-MB-436 ($BRCA1^{-/-}$) cells.**

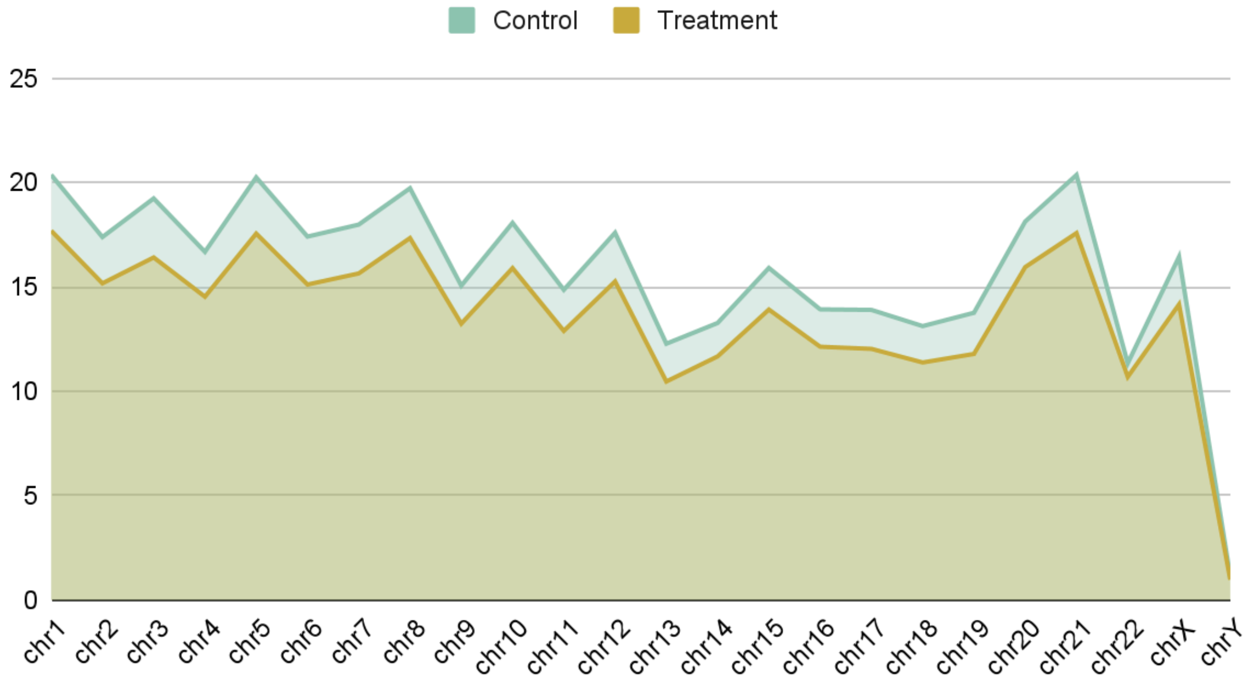


**Supplementary Figure S8. Mean sequencing depth per chromosome of control and treatment (olaparib-treated) samples.**
